# SARS-CoV-2 Spike N-Terminal Domain modulates TMPRSS2-dependent viral entry and fusogenicity

**DOI:** 10.1101/2022.05.07.491004

**Authors:** Bo Meng, Rawlings Datir, Jinwook Choi, CITIID-NIHR BioResource COVID-19 Collaboration, John Bradley, Kenneth GC Smith, Joo Hyeon Lee, Ravindra K. Gupta

## Abstract

Over 20 mutations have been identified in the N-Terminal Domain (NTD) of SARS-CoV-2 spike and yet few of them are fully characterised. Here we first examined the contribution of the NTD to infection and cell-cell fusion by constructing different VOC-based chimeric spikes bearing B.1617 lineage (Delta and Kappa variants) NTDs and generating spike pseudotyped lentivirus (PV). We found the Delta NTD on a Kappa or WT background increased spike S1/S2 cleavage efficiency and virus entry, specifically in Calu-3 lung cells and airway organoids, through use of TMPRSS2. We have previously shown Delta spike confers rapid cell-cell fusion kinetics; here we show that increased fusogenicity can be conferred to WT and Kappa variant spikes by transfer of the Delta NTD. Moving to contemporary variants, we found that BA.2 had higher entry efficiency in a range of cell types as compared to BA.1. BA.2 showed higher fusogenic activity than BA.1, but the BA.2 NTD could not confer higher fusion to BA.1 spike. There was low efficiency of TMPRSS2 usage by both BA.1 and BA.2, and chimeras of Omicron BA.1 and BA.2 spikes with a Delta NTD did not result in more efficient use of TMRPSS2 or cell-cell fusogenicity. We conclude that the NTD allosterically modulates S1/S2 cleavage and spike-mediated functions such as entry and cell-cell fusion in a spike context dependent manner, and allosteric interactions may be lost when combining regions from more distantly related spike proteins. These data may explain the lack of successful SARS-CoV-2 inter-variant recombinants bearing breakpoints within spike.

## Introduction

The SARS-CoV-2 furin cleavage site (also known as a polybasic cleavage site), cleaved in producer cells by a ubiquitously expressed protease furin (Hoffmann et al., 2020a), is believed to be one of the main reasons behind the success of SARS-CoV2 worldwide (Johnson et al., 2021; Peacock et al., 2021). Upon virus release the trimeric spike engages with angiotensin converting enzyme 2 (ACE2) of the target cells to initiate virus entry (Jackson et al., 2022; Peng et al., 2021). Depending on the abundance of the cofactor TMPRSS2 and the status of spike cleavage (Hoffmann et al., 2020b; Ou et al., 2021a, 2021b; Park et al., 2016; Whittaker et al., 2021), the virus either enters through the TMPRSS2 mediated route by fusing at the plasma membrane or via late endosomes through a secondary cleavage preceding the concealed fusion peptide at S2’ (Figure 1A). The latter does not require a pre-cleaved spike at S1/S2, as cathepsin can mediate cleavage of both S1/S2 and S2’.

**Figure 1:**
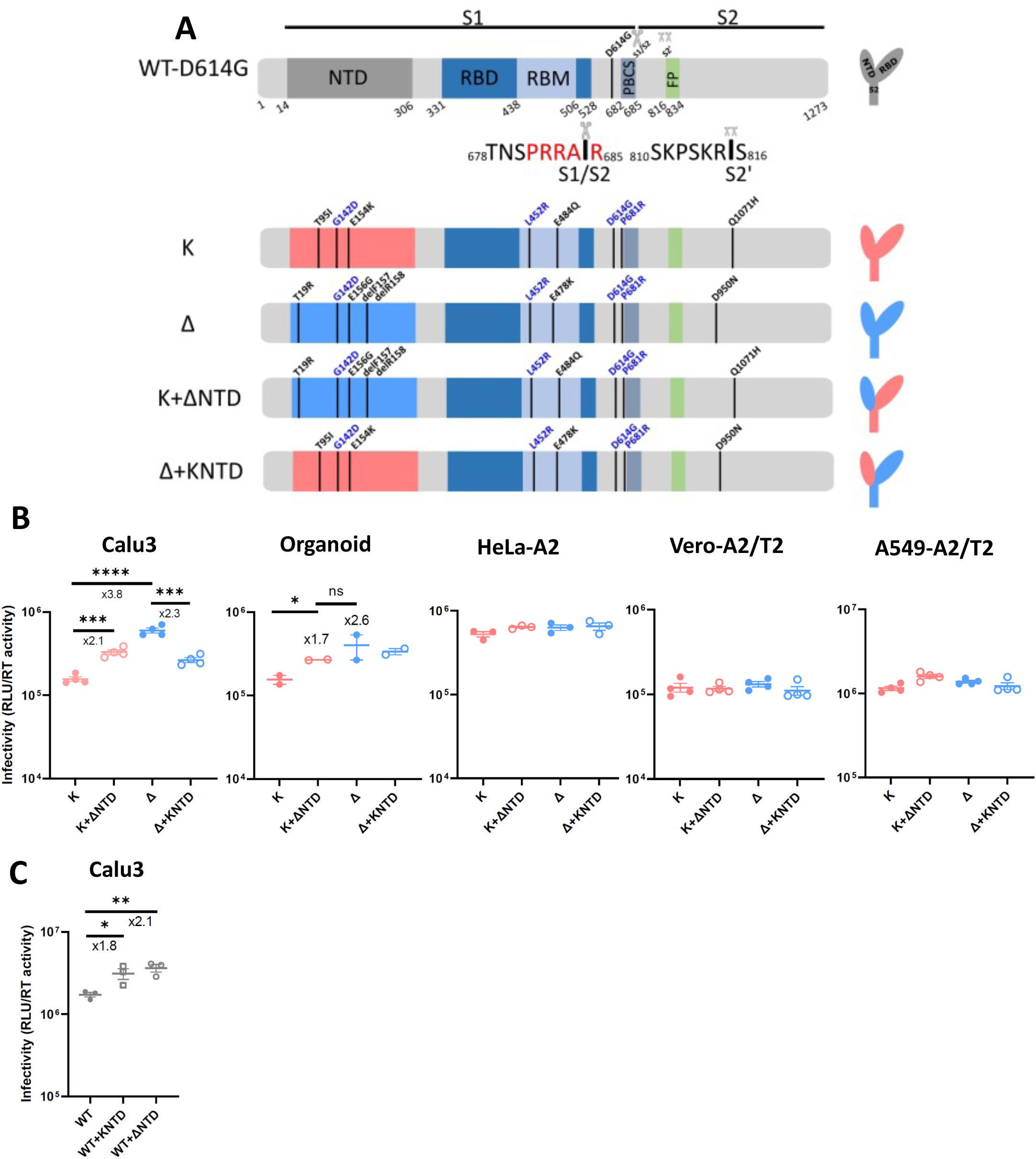
Delta exhibits increased infectivity over Kappa in Calu3 cells and is dependent on the NTD. (A): Schematic diagrams of WT, Kappa and Delta with their chimeras bearing swapped NTD. The consensus mutations between Kappa and Delta are annotated in blue. The monomeric spikes shown on the right-hand side are for illustration purposes. (B): PV bearing Delta or Kappa or chimeric spike was used to transduce Calu3 and organoids expressing endogenous levels of ACE2 and TMPRSS2 and ACE2/TMPRSS2 overexpressing cell lines including Hela-ACE2, Vero-ACE2/TMPRSS2 and A549-ACE2/TMPRSS2. Unpaired t test. (C): WT, WT with KappaNTD and WT with DeltaNTD were used to transduce Calu3 cells. In B&C, mean +/- SEM are shown for technical replicates (n=2-4). Two-sided unpaired Student t test, ns: not significant, *p<0.05, **p<0.01, ***p<0.001, ****p<0.0001. Data are representative of at least two experiments.

Kappa (B.1.617.1) and Delta (B.1.617.2) SARS-CoV-2 variants were first detected in India in late 2020 (Dhar et al., 2021; Ferreira et al., 2021). Although Kappa, the earliest detected B.1.617 variant in India, displayed greater escape to vaccine-elicited antibody responses (McCallum et al., 2021a), Delta surpassed Kappa to become the dominant strain in India and worldwide by mid-2021 (Mlcochova et al., 2021). The S gene encodes 8 and 9 non-synonymous mutations in Kappa and Delta, respectively and four of which are shared by both (Figure 1A). The NTD is more variable between the two (3 and 4 mutations relative to Wu-1 in Kappa and Delta, respectively). In contrast, the RBD, each bearing two mutations, is more constrained most likely due to its obligatory role in engaging with ACE2. Diverse mutations in the NTD are not unique to Kappa or Delta and have been documented in Alpha and many other VOCs (McCarthy et al., 2021). Additionally, both Kappa and Delta share two conserved mutations at D614G and P681R and a unique mutation at S2; Q1071H for Kappa and D950N for Delta.

We previously showed that both Kappa and Delta spikes exhibit highly efficient cleavage of S1/S2 over 614G WT (Mlcochova et al., 2021). Intriguingly Delta appears to be superior to Kappa in entering Calu3 cells and organoids expressing endogenous level of ACE2 and TMPRSS2. Receptor binding is modestly increased in Delta but lower than the preceding strains of VOCs, for example Alpha (Collier et al., 2021a; Liu et al., 2022; Ulrich et al., 2022), indicating enhanced receptor binding cannot solely explain the higher transmissibility of Delta (Barton et al., 2021; McCallum et al., 2021a; Ramanathan et al., 2021; Saville et al., 2022; Supasa et al., 2021). Moreover, cryo-EM structures of Delta and Kappa trimeric spike show that both RBD and S2 adopt a remarkably similar geometry suggesting those two regions are less likely to be accountable for the increased entry of Delta over Kappa (Zhang et al., 2021a). *In vitro*studies using replication competent virus isolates showed that Delta has fast replication kinetics in Calu3, human airway epithelium cells and in airway organoids (Mlcochova et al., 2021). However, the underlying molecular mechanism for the high transmissibility of Delta over Kappa in the real world is elusive.

Our published data also showed that the RBD on its own did not confer higher infectivity to Kappa (Ferreira et al., 2021), suggesting the NTD may be responsible for the increased infectivity. The NTD interacts with cofactors L-SIGN and DC-SIGN at the cell surface (Lempp et al., 2021); blockade of these proteins can effectively neutralise the virus in ACE2 non-overexpressing cells, suggesting NTD and RBD may work cooperatively. The cooperativity of NTD and the RBD is additionally supported by the identification of infectivity enhancing antibodies specifically targeting the NTD domain (Li et al., 2021; Liu et al., 2021), and the observation that binding of the 4A8 monoclonal antibody in the NTD modulates the RBD into an up position (Chi et al., 2020; Díaz-Salinas et al., 2022). Interestingly such antibody binding sites coincide with known infectivity enhancing sites, such as the H69V70 deletion that emerged during an example of intra-host evolution (Kemp et al., 2021) and in Alpha (Meng et al., 2021) and Omicron variants (Meng et al., 2022). It is therefore plausible that the NTD plays an active role in virus entry by engaging with host cofactors and triggering conformational changes of the RBD.

Despite over 20 mutations documented in the NTD, the role of those mutations in infectivity and their impact on the immune response elicited by vaccines is less clear. We reported that the H69V70 deletion found in Alpha was positively selected to increase its infectivity with a modest decrease in immune evasion (Meng et al., 2021). Here we hypothesised that the NTD plays a regulatory role that impacts S1/S2 cleavage and ACE2 usage. We constructed a panel of chimeric spike proteins with NTD from different VOCs in a variety of VOC backbones. We examined those chimeras alongside the parental VOCs in pseudovirus based entry assays (Mlcochova et al., 2020) and investigated spike-mediated fusogenicity. Our data are consistent with a model whereby the NTD regulates virus entry and cell-cell fusion in a variant context-dependent manner.

## Results

### NTD increases Delta infectivity in lung cells and airway organoids

The most dramatic changes in spike between Kappa and Delta lie in the NTD. Both Kappa and Delta spike are efficiently cleaved in the producer cells (Mlcochova et al., 2021). We sought to assess the contribution of the NTD in spike cleavage in purified PV by western blot. We included a deletion mutant in the NTD (delH69/V70) as a control due to its known efficient spike cleavage (Kemp et al., 2021; Meng et al., 2021). Plasmids encoding HIV Gag/pol, a genome flanked by LTRs encoding luciferase, as well as the corresponding spike were transfected into 293T producer cells. The supernatants were harvested and pelleted through ultracentrifugation for western blot analysis. Our data show that the H69/V70 deletion increased S1/S2 cleavage compared to Wu-1 as expected (Figure S1A). Kappa and Delta spike were efficiently cleaved with a more pronounced cleavage observed in Delta (Figure S1B). We additionally observed that the Kappa spike was prone to S1 shedding evidenced by a higher S2/S1 ratio compared to that of the WT (Figure S1C). In contrast, Delta spikes were more stable. We further noticed that the increased efficiency in spike cleavage requires a cognate NTD as the spike bearing the RBD mutations alone failed to be cleaved efficiently (Figure S1D), suggesting NTD on its own or together with the RBD influences the cleavage activity. We conclude that the Delta spike has evolved to be optimal in efficient spike cleavage whilst maintaining spike stability, reminiscent of the emergence of D614G in the early pandemic (Gobeil et al., 2021; Yurkovetskiy et al., 2020; Zhang et al., 2021b, 2020).

We previously observed enhanced cell-free infectivity for Delta over Kappa in Calu3 cells in the PV system (Mlcochova et al., 2021). Importantly this difference was also evident for airway organoids (Mlcochova et al., 2021; Youk et al., 2020) suggesting that at endogenous expression levels of ACE2 and TMPRSS2, Delta has enhanced ability over Kappa in virus entry. Calu-3 and airway organoids express high levels of TMPRSS2 (Meng et al., 2022). Given that cleavage efficiency positively correlates with the infectivity in lung cells (Meng et al., 2022), we sought to examine whether the Delta NTD also contributes to higher infectivity over Kappa in vitro.

To test this we constructed chimeras harboring either the Delta NTD or the Kappa NTD in Delta or Kappa backbones (Figure 1A). We then used the PV bearing different spikes to transduce cell lines to examine the entry efficiency. Interestingly we observed elevated cell-free infectivity in a Kappa chimera bearing the Delta NTD in Calu3 cells and primary human airway organoids but not in other cell lines, indicating that the NTD confers the specificity for this increase (Figure 1B). Consistent with this model, a decrease in infectivity in Calu3 cells and primary human airway organoids was observed when the Kappa NTD was exchanged with that of Delta. We further extended this observation to the WT backbone where a hybrid WT spike bearing either Kappa or Delta NTD was expressed (Figure 1C and S2). Consistent with these observations, both Kappa and Delta NTD when fused with WT increased cell-free infectivity in Calu3 lung cells.

### The NTD influences virus entry efficiency

We next sought to delineate the underlying mechanism for this increased infectivity in Delta over Kappa. Spike cleavage correlates with SARS-CoV-2 virus entry pathway preference (Meng et al., 2022; V’kovski et al., 2021). Western blots using the cell lysates from the 293T virus producer cells showed that the expression levels of chimeric spikes were comparable (Figure 2A&B). However, the cleavage of spike in the chimeras phenocopied the cleavage patterns from which their NTD were derived (Figure 2C). We extended this observation in WT chimeras where increased spike cleavage was also evident following addition of either Kappa NTD or Delta NTD (Figure S2B). Structural data indicate that the NTD, RBD and PBCS are cooperatively regulated (Gobeil et al., 2021). Hence we reasoned that the observed increase in the accessibility to the furin cleavage site may be regulated by the allostery involving the NTD.

**Figure 2:**
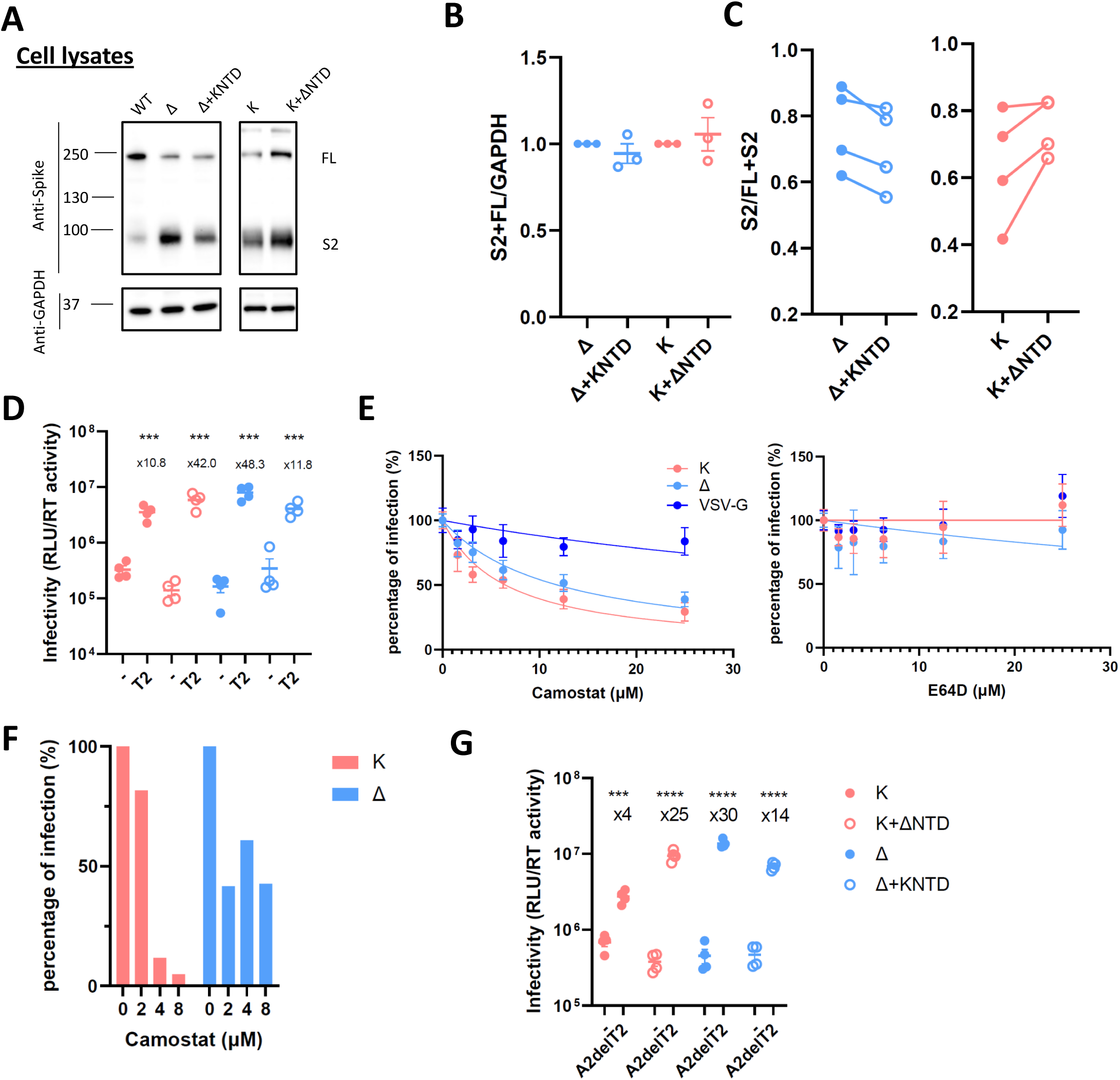
The NTD modulates the usage of TMPRSS2 and ACE2 for virus entry. (A): A representative western blot of cell lysates showing spike cleavage in Delta, Kappa and its chimeras. GAPDH was probed as a loading control. (B): The band intensity from (A) was densitometrically calculated using ImageJ. Total S2 associated spike proteins (S2 and FL) were then normalized against GAPDH across Delta, Kappa and NTD bearing chimeras. (C): The ratio of S2/FL+S2 from the band intensity of S2 and FL shown in (A) was plotted. n≥3. (D): Delta, Kappa or chimeras was transduced into either parental 293 cells or 293T cells overexpressing TMPRSS2. The fold increase of the virus entry in T2 overexpressing cells over parental cells is shown above the scatter plots. mean +/- SEM are shown for technical replicates (n=4). Two-sided unpaired Student t test, ***p<0.001. Data are representative of two experiments. (E): The entry efficiency of Delta and Kappa in A549-ACE2/TMPRSS2 cells in the presence of TMPRSS2 specific inhibitor camostat or cathepsin specific inhibitor E64D. The lentivirus pseudotyped with VSV-G was used as a control. The RLU was normalized with non-drug added control giving a percentage of infection. The data showing the mean±SD from 4 experiments were plotted. (F): The entry efficiency of Delta and Kappa in the presence of camostat in airway organoids. Data are representative of two experiments. (G): Delta, Kappa or chimeras was transduced into either parental 293T cells or 293T cells overexpressing ACE2 with abrogated TMPRSS2 expression (A2delT2). The fold increase of the virus entry in A2delT2 overexpressing cells over parental cells is shown above the scatter plots. mean +/- SEM are shown for technical replicates (n=4). Two-sided unpaired Student t test, ***p<0.001, ****p<0.0001. Data are representative of two experiments.

SARS-CoV-2 primarily enters Calu-3 cells through membrane fusion due to an abundant expression of TMPRSS2 and ACE2 at the plasma membrane. The primary cleavage at S1/S2 is a prerequisite for a secondary cleavage at S2’ site by proteases such as TMPRSS2 at the plasma membrane (V’kovski et al., 2021), triggering the conformational rearrangement of S2 for the exposure of the fusion peptide. Due to more efficient cleavage of the Delta spike (Figure S1), we reasoned that the Delta spike may be more efficiently primed to use TMPRSS2 at the plasma membrane for its entry. To test this hypothesis, we transduced the virus bearing either authentic spike of Delta or Kappa or their NTD-chimeric counterparts into TMPRSS2 overexpressing 293T cells or parental 293T cells (Figure 2D). As expected, all the viruses including chimeras increased entry efficiency when TMPRSS2 was present with an even more pronounced increase observed in Delta, consistent with the observation that Delta is more efficient in utilising TMPRSS2. Intriguingly, we also observed an increased transduction efficiency for the Kappa chimera that was comparable to that of Delta, suggesting that the Delta NTD enables the entry of chimeric Kappa by efficient TMPRSS2 usage. Consistent with this model, the ratio of entry for the Delta chimera containing Kappa NTD entry in TMPRSS2 overexpressing cells versus normal expressing cells was reduced.

To further investigate the dependence of TMPRSS2 on virus entry, we pretreated A549-ACE2/TMPRSS2 cells or airway organoids with either camostat (a TMPRSS2 inhibitor) or E64D (a cathepsin inhibitor). We observed that the IC50 of camostat in Delta was approximately 2 fold higher than Kappa, whilst E46D had little effect on either virus (Figure 2E). Reassuringly, a similar observation was made in airway organoids (Figure 2F), suggesting that Delta is more resistant to camostat and hence is more efficient in utilising available TMPRSS2 for virus entry. Indeed the Kappa chimera bearing Delta NTD showed a similar drug sensitivity profile to Delta implying that the NTD is accountable for this shift in TMPRSS2 sensitivity (Figure 3S).

**Figure 3:**
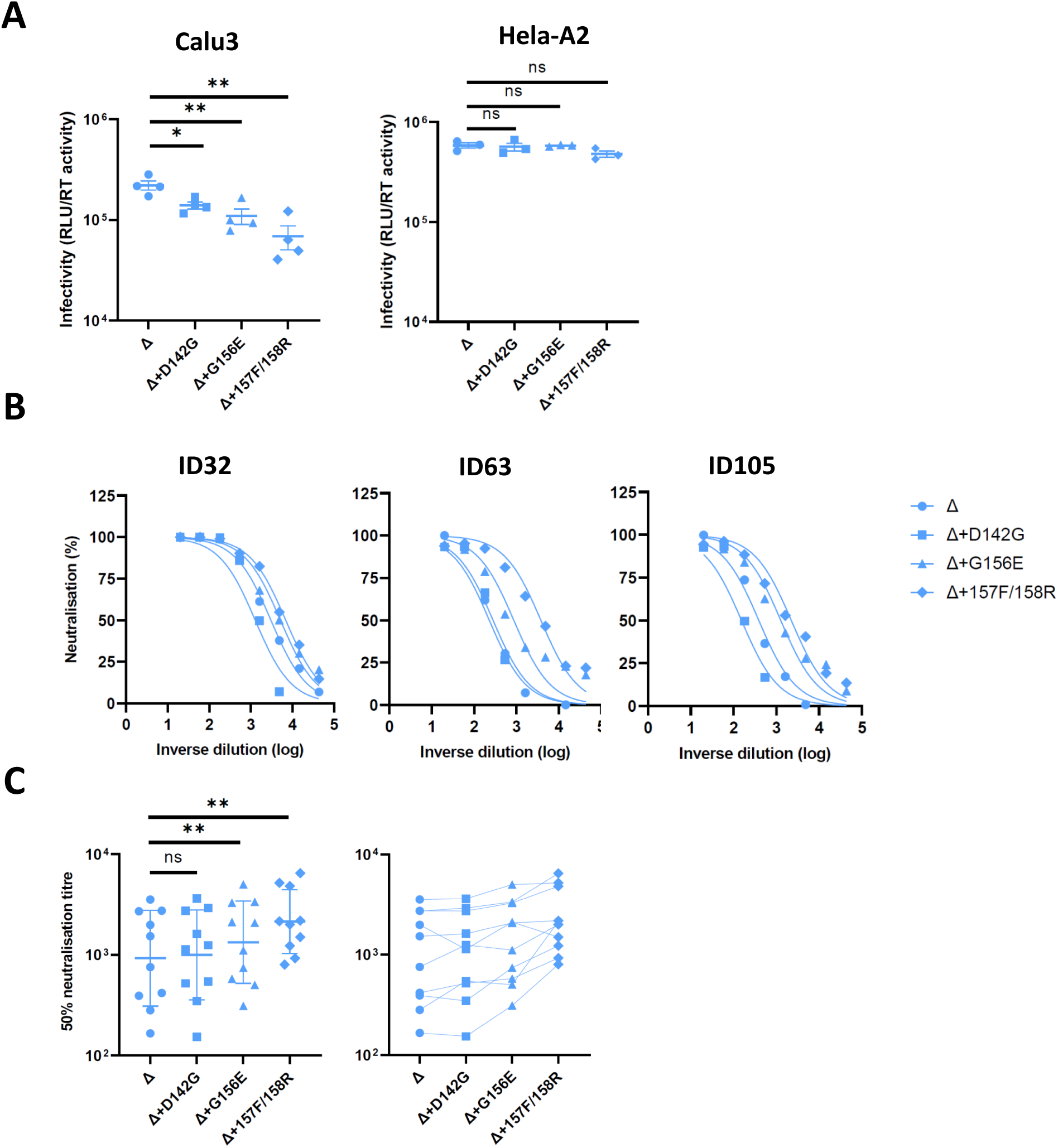
Reversing the mutations in the Delta NTD reduces the infectivity in lung cells and increases the neutralisation sensitivity to the vaccine-elicited antibody. (A): PV bearing Delta and its reversions were used to transduce Calu3 and Hela-ACE2 cells. mean +/- SEM are shown for technical replicates (n=4). Two-sided unpaired Student t test. Data are representative of two experiments. (B): Examples of neutralisation curves from ID32, 63 and 105 vaccinees with PVs bearing the reversion at 142, 156 or 157/8. (C): The 50% serum neutralisation was plotted across ten sera showing the geometric mean with geometric SD. Paired Wilcoxon was used for analysis. Data are representative of two experiments. ns: not significant, *p<0.05, **p<0.01.

It is reported that the NTD modulates RBD conformation through allosteric effects (Qing et al., 2021). We speculated the accessibility to ACE2 is altered in the chimeric spikes. We next went on to examine this by transducing parental 293T cells or their isogenic counterparts where the expression of TMPRSS2 is abrogated and ACE2 is overexpressed (A2delT2). Indeed, we found a 30 fold increase in Delta when ACE2 is overexpressed whereas Kappa only had 4 fold increase (Figure 4G). More strikingly, the Kappa chimera, bearing the Delta NTD in the Kappa spike backbone, showed a similar magnitude of increase as Delta. Conversely, the Delta chimera bearing a Kappa NTD manifested a decrease in ACE2 dependence compared to Delta, albeit to a lesser extent. Taken together, our data support the notion that the Delta NTD modulates the use of TMPRSS2 more efficiently due to spike cleavage and that the Delta NTD allosterically regulates the RBD to increase efficiency of ACE2 usage.

**Figure 4:**
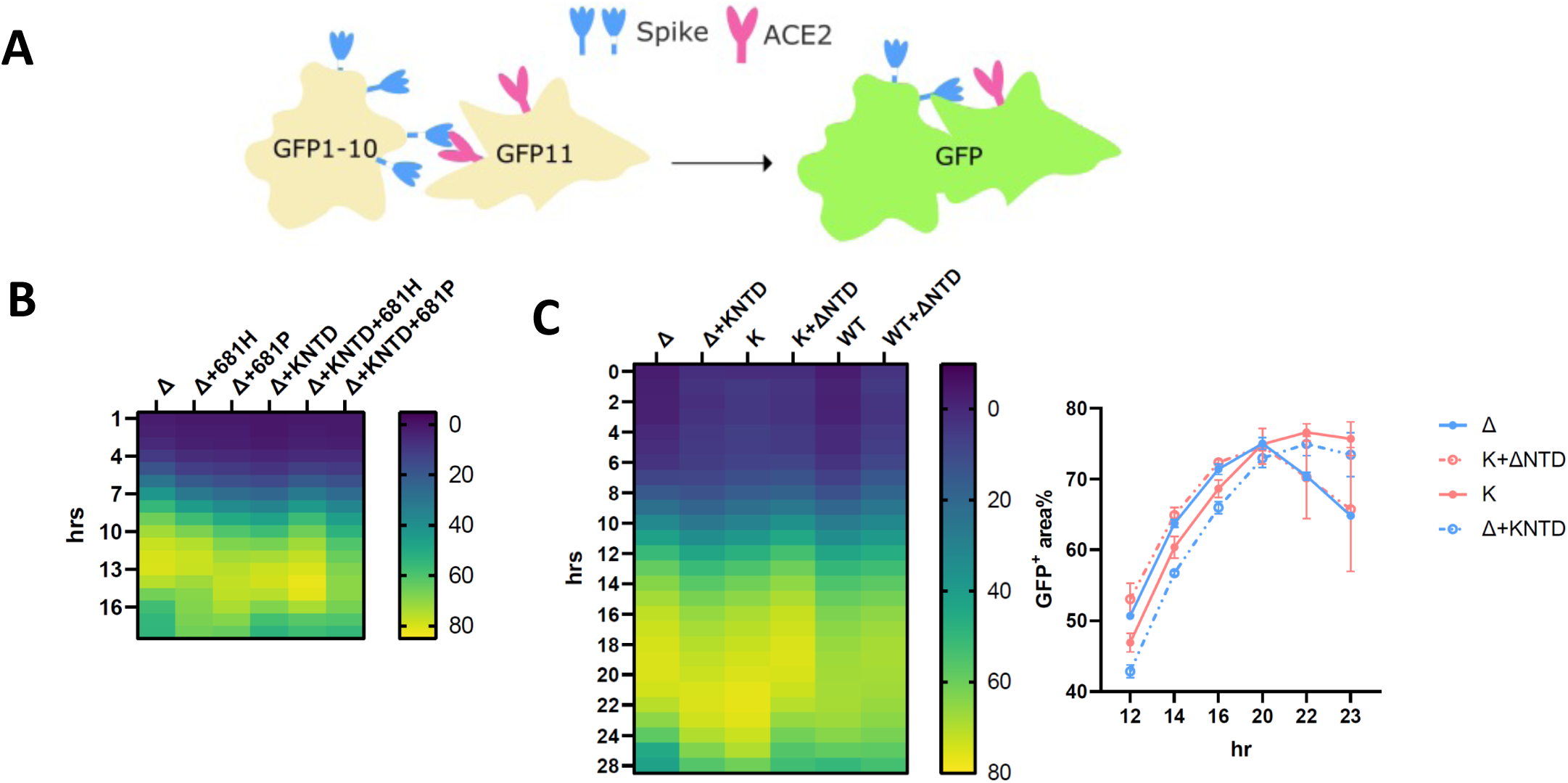
The SARS-CoV-2 Delta NTD increases the fusion kinetics of Kappa and WT spikes. (A): A schematic diagram showing the split GFP system for spike-ACE2 mediated cell fusion. (B): 681R or 681H is required for the enhanced fusogenicity in Delta and its chimera bearing Kappa NTD. (C): The fused Delta NTD in Kappa and WT increased the fusion kinetics of their counterparts, respectively. The line graphs on the right show the percentage of the positive GFP area at 12, 14, 16, 20, 22 and 23 hr post transfection. The data showing the SEM at each point were averaged from two experiments. The heatmap at each time point shows the mean of the GFP positive area over the field of view from two experiments.

### NTD mutations and deletions impact entry efficiency

To probe the mutations in the Delta NTD that may contribute to the enhanced infectivity in Calu3 cells, we constructed a series of spike bearing PV by reverting a cluster of amino acids (142D, 156G, del157F and del158R) to WT Wuhan-1 residues individually. Interestingly, each individually reverted mutant showed a decrease in infectivity in Calu3 cells with the greatest reduction seen following re-insertion of the deleted 157R/158R (3 fold). This is in agreement with the infectivity reduction in the Delta NTD bearing Kappa chimera (Figure 1B). The decrease in infectivity was only observed in Calu3 cells but not in Hela-ACE2 cells, consistent with the cell specificity in virus entry conferred by the NTD. Since the NTD harbors the antigenic supersites that are mutated in Delta (Cerutti et al., 2021; McCallum et al., 2021b; McCarthy et al., 2021; Suryadevara et al., 2021), we predicted that the reversion of such mutations would at least partially rescue the loss of sensitivity to the vaccine sera. To test this, PV were harvested and used to transduce HeLa-ACE2 cells in the presence of a dilution series of vaccine sera from vaccinees who had received two doses of BNT162b2 at least 1 month prior to sampling. D142G alone did not alter the sensitivity of neutralisation (Figure 3). However, G156E or repairing the deletion of 157F/158R increased the sensitivity of neutralization by 2 fold, consistent with the notion that the NTD is important for immune evasion and infectivity.

### Delta NTD confers faster cell-cell fusion kinetics in a context-dependent manner

Syncytium formation, mediated by spike and the host receptor ACE2, has been found in SARS-CoV-2 infected patients and is thought to be important for disease progression (Braga et al., 2021). We and others had previously demonstrated that variants bearing furin cleavage site mutations exhibit a pronounced increase in fusogenicity, with the exceptions of Omicron (Figure 4A) (Buchrieser et al., 2021; Meng et al., 2021, 2022; Mlcochova et al., 2021; Rajah et al., 2021). Given that spike cleavage largely correlates with fusion and we observed an enhanced cleavage in the Kappa chimera, we sought to examine whether spike mediated fusion is altered. We first confirmed that efficient cleavage is required for the syncytia formation as the mutants bearing the WT proline at position 681 (681P) decreased fusion efficiency in both parental and chimeras whereas the mutation to histidine (681H), the other cleavage site mutation firstly found in Alpha and then in Omicron, did not (Figure 4B). We then went on to explore whether or not the fusogenicity is different between Delta and Kappa and their chimeras. Concordant with the published studies we found a marked increase in fusion for Delta and Kappa compared to WT (Figure 4C) (Mlcochova et al., 2021; Rajah et al., 2021). Unexpectedly, we found the fusion kinetics between Delta and Kappa were noticeably different despite manifesting a comparably maximal fusion activity at the steady-state (around 70%; Figure 3C). More specifically, Delta was more fusogenic at earlier time points whereas Kappa required two more hours to reach a similar intensity. Intriguingly when the Kappa NTD was fused into Delta the fusion phenotype shifted from Delta to Kappa, with a slowing of kinetics (Figure 3C). Conversely, we observed a fast-fusing Delta phenotype when the Delta NTD was swapped into Kappa. Of note, we also observed a faster fusion phenotype when the Delta NTD was fused into the WT backbone, though this effect was less pronounced compared to that observed in the Kappa chimera (Figure 4C). Taken together we have demonstrated that the Delta NTD can drive faster fusion in both WT and Kappa spike backgrounds.

Epidemiological studies suggest that the more recent Omicron (BA.1 and BA.2) variants are distantly related to all the other early pandemic variants (Simon-Loriere and Schwartz, 2022). It therefore remains possible that the ability of the Delta NTD to drive faster fusion is dependent on their spike background. We found that in comparison to BA.1 BA.2 was more efficient in entering Calu3 cells, though still not as efficient as Delta (Figure 5A). An elevated level of entry of BA.2 over BA.1 was observed in H1299 cells where the entry of Delta was the least efficient. These data suggest that BA.2 is capable of infecting both TMPRSS2 low (H1299) and high (Calu3) cells efficiently.

**Figure 5:**
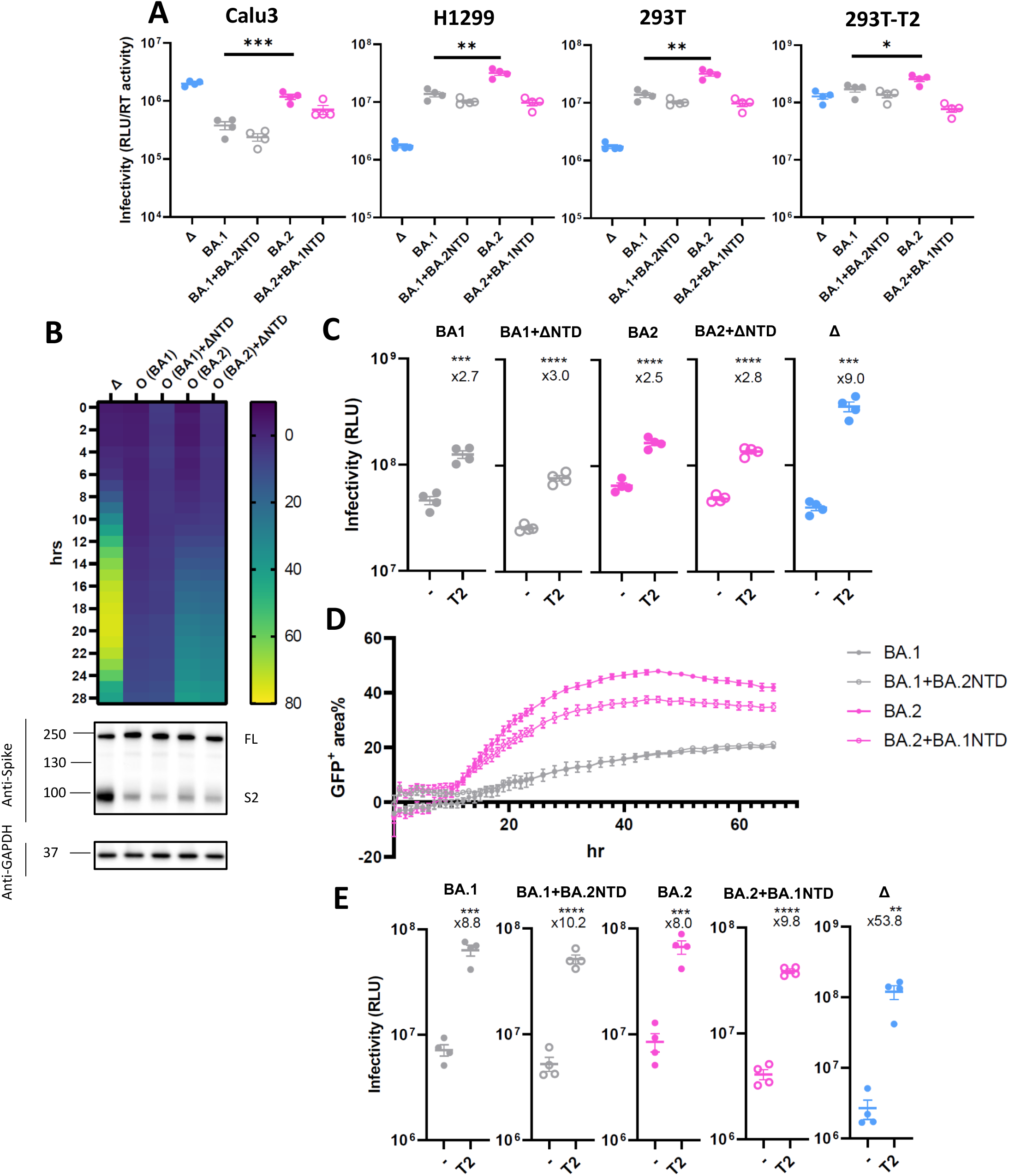
The SARS-CoV-2 Delta NTD or BA.2 NTD does not alter spike fusion or sensitivity to TMPRSS2 of BA.1. (A): PV bearing Delta, BA.1, BA.2 or chimeric forms of BA.1 and BA.2 spike were used to transduce Calu3, H1299 and 293T expressing endogenous levels of ACE2 and TMPRSS2 and TMPRSS2 overexpressing 293T cells. (B): The fusion kinetics of the fused Delta NTD in BA.1 and BA.2 along with their parental spikes. The western blot showing cleavage of spike is directly underneath the heatmap. (C): BA.1, BA.2 or chimeras with Delta were transduced into either parental 293 cells or 293T cells overexpressing TMPRSS2. The fold increase of the virus entry in TMPRSS2 overexpressing cells over parental cells is shown above the scatter plots. (D): The fusion assay of BA.1, BA.2 and their chimeric BA.1 bearing BA.2 NTD and BA.2 bearing BA.1 NTD. The line graphs show the percentage of the positive GFP area at 1 hr interval post transfection. The data showing the SEM at each point were averaged from two experiments. (E): BA.1, BA.2 or chimeras bearing BA.2 NTD or BA.1 NTD together with Delta were transduced into either parental 293 cells or 293T cells overexpressing TMPRSS2. The fold increase of the virus entry in T2 overexpressing cells over parental cells is shown above the scatter plots. In A, C&E, the plots are representative from two experiments. mean +/- SEM are shown for technical replicates (n=4). Two-sided unpaired Student t test, *p<0.05, **p<0.01, ***p<0.001, ****p<0.0001.

We then introduced the Delta NTD into the both BA.1 and BA.2 to examine whether the Delta NTD could also confer increased fusion to Omicron, a known poor cell-cell fusion spike (Figure 5B). Although BA.2 showed faster fusion than BA.1, consistent with a recently published study (Yamasoba et al., 2022), both Omicron and the Delta-NTD fused Omicron chimeras showed poor spike-mediated fusion and that most importantly the difference between Omicron and its counterpart was negligible. Intriguingly, this phenotype correlates with inefficient cleavage of spike even after the cleavage-enhancing Delta NTD was fused into an Omicron spike background.

Omicron has evolved to avoid the early entry route via TMPRSS2 at the plasma membrane by entering the cells through endosomal route (Meng et al., 2022). Given spike cleavage was not affected by domain swapping, we predicted that the Omicron chimera bearing the Delta NTD would fail to confer increased sensitivity to TMPRSS2, in contrast to our findings for the B.1.617 lineages Delta and Kappa. Indeed there was only a moderate increase in entry (2-3 fold) when TMPRSS2 was overexpressed, compared to 9 fold for Delta, reinforcing the notion that the virus entry of Omicron is TMPRSS2 independent and that altering the NTD is not sufficient to confer TMPRSS2 usage to Omicron.

We next sought to examine whether the BA.2 NTD could rescue the poor fusogenicity of BA.1 and/or whether the BA.1 NTD impaired the fusogenicity of BA.2. In contrast to the fast fusion activity conferred by the Delta NTD in Kappa or WT (Figure 4C), BA.2 NTD failed to increase the fusion kinetics of the BA.1 chimeric spike bearing BA.2 NTD (Figure 5D). The BA.1 NTD modestly suppressed the fusion of BA.2 chimeric spike bearing BA.1 NTD. Interestingly, when the sensitivity to TMPRSS2 was probed (Figure 5E), we found little difference between BA.1 and BA.2 and their chimeras (8-10 fold) in comparison to Delta (over 50 fold), indicating that in Omicron the NTD on its own is not sufficient to affect TMPRSS2 utilisation. Taken together, these data strongly suggest that the enhanced fusogenicity of the Delta NTD and the spike cleavage is context-dependent and that non-NTD regions may be involved to suppress efficient fusion activity of Omicron spike.

## Discussion

Cooperativity between the NTD and RBD is reflected by the finding that NTD deletions were previously discovered after the emergence of the RBD mutations; for example, N439K and Y453F neutralising antibody escape mutations were identified first before the acquisition of delH69/V70 in the NTD (Meng et al., 2021). We previously showed that some escape mutations reduced virus entry efficiency and the delH69/V70 NTD deletion restored entry whilst maintaining immune evasion (Kemp et al., 2021; Meng et al., 2021). Therefore, viruses have selected various mutations in the RBD due to the immune pressure imposed on this immunodominant site by the host. However these RBD-mutated viruses have suboptimal fitness, thus placing selection pressure to acquire compensatory mutations in the NTD. We propose that NTDs in VOC have evolved under such pressure.

The NTD has been proposed to modulate the RBD to a receptor accessible mode (Qing et al., 2021). We speculated that by alternating the NTD of Delta into Kappa the receptor engagement with ACE2 is affected. Indeed we observed an enhanced virus entry in the Kappa chimera bearing the Delta NTD (Figure 2G). However an increased accessibility to ACE2 cannot solely account for the enhanced infectivity observed in Calu3 as such an increase is not evident for the cell lines where ACE2 is also abundantly expressed. A marked increase for chimeric Kappa spike to use TMPRSS2 at the plasma membrane provides a plausible explanation for this specificity; the chimeric Kappa spike is as efficient as Delta on TMPRSS2 usage (Figure 2D). This observation is in agreement with our inhibitor experiment, whereby the addition of the TMPRSS2 protease inhibitor camostat has a more pronounced inhibitory effect for Kappa than Delta (Figure 2E&F). However, the Delta NTD cannot enhance the cleavage of Omicron spike nor its dependence on TMPRSS2 (Figure 5B&C), suggesting that the presence of the Delta NTD is not the sole determinant of TMPRSS2 usage. Furthermore, although allosteric conformational changes involving NTD and other regions may contribute to phenotype, binding of the NTD to a secondary cofactor at the plasma membrane cannot be completely excluded (McCallum et al., 2021b).

The NTD of VOCs, including poorly fusogenic Omicron (Meng et al., 2022; Suzuki et al., 2022), once fused with WT exhibits an increased fusion in the presence of trypsin indicating the accessibility of spike cleavage by host protease is affected through the allostery of the NTD (Qing et al., 2022). We have proposed that the sensitivity to TMPRSS2 correlates with cleavage status at S1/S2, which has an additional impact on the cell to cell fusion (Meng et al., 2022). Consistent with this notion, we found the Delta NTD confers faster fusion kinetics in both WT and Kappa which coincides with an elevated S1/S2 cleavage in WT and Kappa spike bearing the Delta NTD. However S1/S2 cleavage is unaltered in the Delta NTD bearing Omicron which also has poor fusogenicity. These data demonstrated that regions other than NTD may also be required for efficient cleavage. Indeed a recent study using a domain swapping approach has suggested the RBD of Omicron contributes to inefficient spike cleavage and inferior fusogenicity (doi.org/10.1101/2022.04.03.486864). The specific site in S2 preceding to fusion peptide has also been implicated in facilitating an efficient cleavage in accordance with the NTD (Qing et al., 2022).

We observed that the Kappa spike is less stable than that of Delta (Figure S1C). We suspect that when Delta first emerged with the RBD mutations the intermediate strain was less fit leading to a premature S1 shedding. However, the subsequently acquired mutations in the NTD allosterically foster non-covalent interactions between S1 and S2 forming more stabilised protomers before engagement with ACE2. This process ultimately leads to active S1 shedding and exposure of the fusion peptide. D614G emerged as a positively selected mutation in an early stage of the pandemic persisting in subsequent lineages by stabilising trimeric spike. The more stable form of S1 is therefore better positioned to engage with ACE2 (Daniloski et al., 2021; Díaz-Salinas et al., 2022; Yurkovetskiy et al., 2020; Zhang et al., 2021b, 2020). It is reasonable to speculate that the effect of the NTD mutations is analogous to the acquisition of D614G by strengthening the intramolecular interactions to prevent S1 from being loosened prematurely while bolstering the RBD in an ACE2-accessible form.

Finally we showed that Delta NTD revertant mutations (G156E and 157F/158R repair) were able to compromise cell entry and increase sensitivity to neutralisation by vaccine sera. Reassuringly, a similar finding was recently reported in breakthrough B.1.617 viruses where the same mutations in the NTD accounted for the attenuated neutralisation sensitivity (Mishra et al., 2022).

In summary, our data support the cooperativity between NTD, RBD and the furin cleavage site. We propose that the NTD allosterically affects the conformation of the RBD for ACE2 binding and the furin cleavage site for spike cleavage. The pre-cleaved spike confers specificity on virus entry via TMPRSS2 usage and drives a faster virus spread with more efficient spike-mediated fusion. Our model explains the dominance of Delta over Kappa by being highly immune evasive on the one hand through the acquisition of known escape mutations, while on the other hand gaining S1/S2 stability and increased infectivity in lung cells. Our study highlights the importance of the continuous effort in monitoring both RBD and NTD mutations in order to understand the biology of variants, and possibly explains the lack of highly fit and successful SARS-CoV-2 inter-variant recombinants bearing breakpoints within spike. On the translational/therapeutic side, combination antibody therapy targeting both NTD and RBD might therefore be less prone to resistance than monotherapy or combinations targeting the RBD alone.

## Acknowledgments

RKG is supported by a Wellcome Trust Senior Fellowship in Clinical Science (WT108082AIA). This study was supported by the Cambridge NIHRB Biomedical Research Centre, Addenbrooke’s Charitable Trust and the Rosetrees Trust. We would like to thank Paul Lehner for Calu-3, James Voss for HeLa ACE2, Simon Cook for H1299 and Suzanne Rihn for the A549 cells. We are grateful to Leo James for 293T-ACE2-ΔTMPRSS2, 293T-TMPRSS2, Vero-GFP1-10 and 293T-GFP11 cells and to Guido Papa and Anna Albecka for the help on imaging. The authors acknowledge plasmids from the G2P-UK National Virology consortium funded by MRC/UKRI (grant ref: MR/W005611/1), and thank Tom Peacock and Wendy Barclay. The views expressed are those of the author(s) and not necessarily those of the NIHR.

## Additional information

Requests for materials, raw data and correspondence should be addressed to Ravindra Gupta

## Author contributions

Conceived research and designed study: R.K.G., B.M, K.G.C.S., J.B. designed experiments: H.Y.L; conducted experiments: B.M, J.C, R.D; B.M, R.K.G wrote the manuscript with input from all authors.

## Methods

### Cells and plasmids

Calu-3 (a human lung epithelial cell line; a gift from Paul Lehner) cells were maintained in Eagle’s minimum essential medium containing 10% FBS and 1% PS. Vero-ACE2/TMPRSS2 cells (a gift from Emma Thomson), Hela-ACE2 (a gift from James Voss) and A549-ACE2/TMPRSS2 (a gift from Massimo Palmarini were maintained in Dulbecco’s modified Eagle’s medium (DMEM) containing 10% FBS and 1% PS. 293T (CRL-3216) and its derivative cell lines including 293TACE2ΔTMPRSS2, 293T-TMPRSS2 and 293T-GFP11 have been described previously (Papa et al., 2021). All the 293T cell lines as well as Vero-GFP1-10 were maintained in DMEM with 10% FBS and 1% PS. All cells were regularly tested and are mycoplasma free. Airway epithelial organoids were obtained and maintained as previously described (Meng et al., 2022; Youk et al., 2020). Human distal lung parenchymal tissues were obtained from adult donors with no background lung pathologies from Papworth Hospital Research Tissue Bank (T02233). Airway organoids were cultured in 48-well plate and were passaged every 2 weeks as previously reported (Meng et al., 2022).

pCDNA_SARS_CoV2_D416G_S WT, Delta, Kappa, Omicron BA.1 and Omicron BA.2 plasmids with 19 amino acid deletion at the C-terminus were generated by gene synthesis. For the construction of the chimeras, the region encompassing the NTD was digested with Bsu36I and HindIII (both NEB) before being gel purified and ligated back to the respective backbone that was cut with the same pair of restriction enzymes. Amino acid substitutions in the Delta NTD (D142G, G156E and repair of 157F and 158R) were introduced into the pCDNA_SARS-CoV-2_ D614G_Delta_S plasmid using the QuikChange Lightning Site-Directed Mutagenesis kit, following the manufacturer’s instructions (Agilent Technologies). Sequences were checked by Sanger sequencing.

### Pseudotype virus preparation and infectivity titration

Plasmids encoding the spike protein of SARS-CoV-2_D614G with a C terminal 19 amino acid deletion were used. Viral vectors were prepared by transfection of 293T cells by using Fugene HD transfection reagent (Promega) as described previously (Kemp et al., 2021). In brief, 293T cells were transfected with a mixture of 11ul of Fugene HD, 1μg of pCDNAΔ19 spike, 1ug of p8.91 HIV-1 gag-pol expression vector and 1.5μg of pCSFLW (expressing the firefly luciferase reporter gene with the HIV-1 packaging signal). Viral supernatant was collected at 48 h after transfection, filtered through 0.45um filter and stored at -80°C. Infectivity was measured by luciferase detection (Bright Glo; Promega) in target cells. The raw readings (in relative light unit (RLU)) were then normalised with the SG-PERT and plotted using GraphPad Prism.

### PV SG-PERT

The SARS-CoV2 spike-pseudotyped viruses containing supernatants were standardised using a SYBR Green-based product-enhanced PCR assay (SG-PERT) as described previously (Pizzato et al., 2009). Briefly, 10-fold dilutions of virus supernatant were lysed in a 1:1 ratio in a 2x lysis solution (made up of 40% glycerol v/v 0.25% Triton X-100 v/v 100mM KCl, RNase inhibitor 0.8 U/ml, TrisHCL 100mM, buffered to pH7.4) for 10 minutes at room temperature. Sample lysates (12 μl) were added to 13 μl of SYBR Green master mix (containing 0.5μM of MS2-RNA Fwd and Rev primers, 3.5pmol/ml of MS2-RNA, and 0.125U/μl of Ribolock RNAse inhibitor and cycled in a QuantStudio (Thermofisher). Relative amounts of reverse transcriptase activity were determined as the rate of transcription of bacteriophage MS2 RNA, with absolute RT activity calculated by comparing the relative amounts of RT to an RT standard of known activity.

### Western blot

For cell lysates, 293 cells were washed and lysed in lysis buffer (Cell Signalling) and lysates were diluted with 4 × sample buffer (Biorad) and boiled for 10 m before subjected to western blotting. For virion purification, clarified supernatants were loaded onto a thin layer of 8.4% optiprep density gradient medium (Sigma-Aldrich) and placed in a TLA55 rotor (Beckman Coulter) for ultracentrifugation for 2 hours at 20,000 rpm. The pellet was then resuspended for western blotting. For protein detection, the following antibodies were used: rabbit anti-SARS-CoV-2 S monoclonal antibody (PA1-41165; Thermofisher), mouse anti-SARS-CoV-2 S1 (MAB105403, R&D systems), rabbit anti-GAPDH polyclonal antibody (10494-1-AP; Proteintech), horseradish peroxidase (HRP)-conjugated anti-rabbit and anti-mouse IgG polyclonal antibody (Cell Signalling). Chemiluminescence was detected using ChemiDoc Touch Imaging System (Bio-Rad). The cleavage ratio of S1 or S2 to FL in virions was determined by densitometry using ImageJ (NIH).

### Drug assay

A549-ACE2-TMPRSS2 (A549-A2T2) cells or human airway organoids were either E64D (Tocris) or camostat (Sigma-Aldrich) treated for 2 hours at each drug concentration before the addition of a comparable amount of input viruses pseudotyped with Delta, Kappa or chimeras (approx. 100,000 RLU). The cells were then left for 48 hours before the addition of substrate for luciferase (Promega) and read on a Glomax plate reader (Promega). The RLU was normalised against the no-drug control which was set as 100%.

### Cell-cell fusion assay

Cell-cell fusion assays were described previously (Meng et al., 2022). Briefly, 293T GFP11 and Vero-GFP1-10 cells were seeded at 80% confluence in a 1:1 ratio in 48 multiwell plate the day before. Cells were co-transfected with 250 ng of spike expression plasmids using Fugene 6 following the manufacturer’s instructions (Promega). Cell-cell fusion was measured using an Incucyte and determined as the proportion of green area to total phase area over time. Data were normalised to non-transfected control. Graphs were generated using Prism 9 software.

### Neutralisation assay

This was performed as previously described (Collier et al., 2021b, 2021a).

---

**Supplementary Figure 1:**
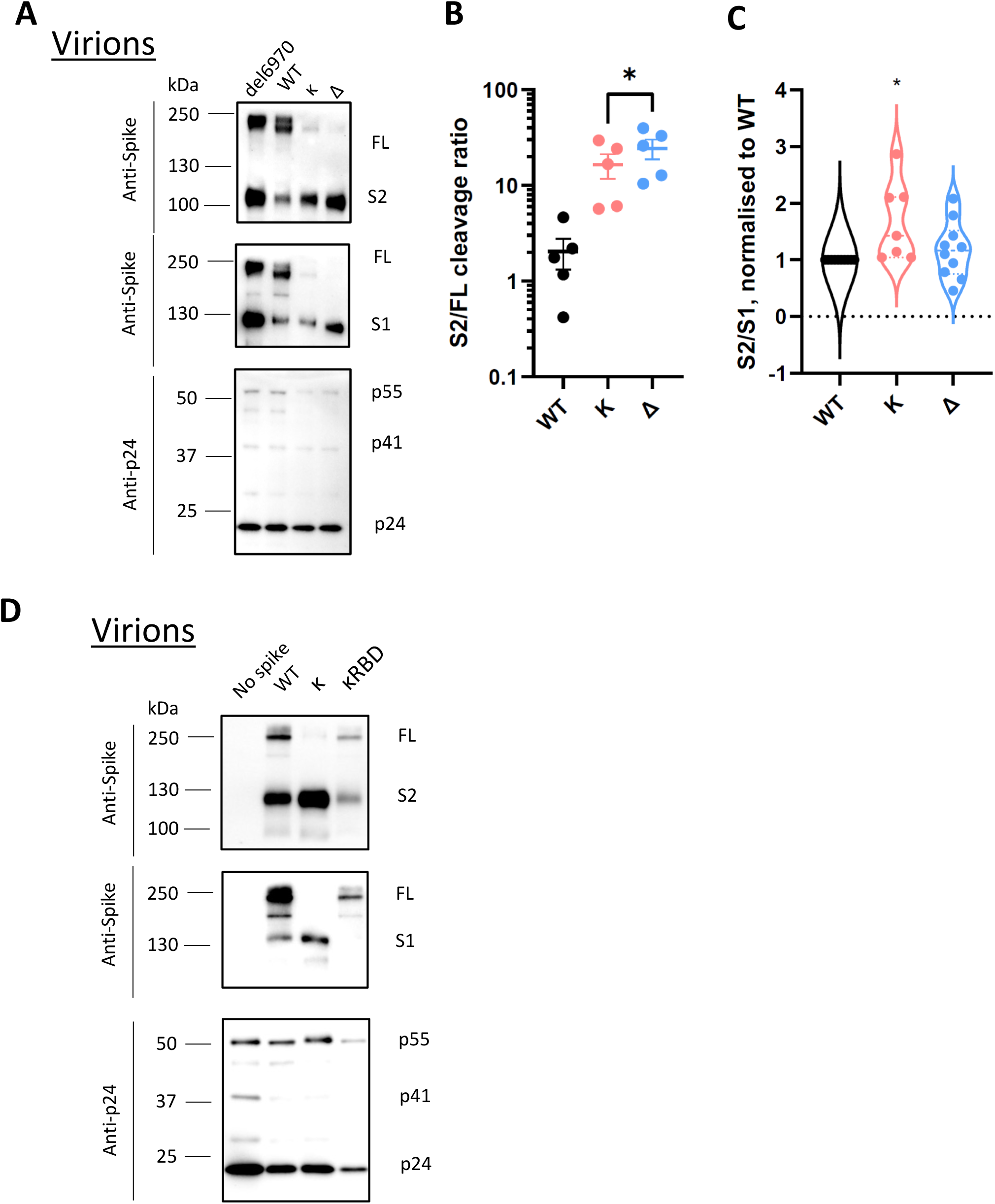
The NTD is required for efficient cleavage of SARS-CoV-2 spike. (A): Western blots of purified PV bearing either H69V70 deletion, WT, Kappa or Delta spikes. The sizes of protein markers were labelled to the left of the blot and the corresponding bands were labelled to the right. (B): The intensity of the spike-associated bands on the western blots was densitometrically quantified (ImageJ) before the ratio was calculated for cleavage (S2/FL, paired t test.) (B) or spike stability (S1/S2; one sample t test) (C). In both (B) and (C) each dot represents one PV preparation. (D): A representative western blot of the purified PVs bearing RBD mutations only in Kappa and Kappa spike, together with WT and non-spike control. *p<0.05.

**Supplementary Figure 2:**
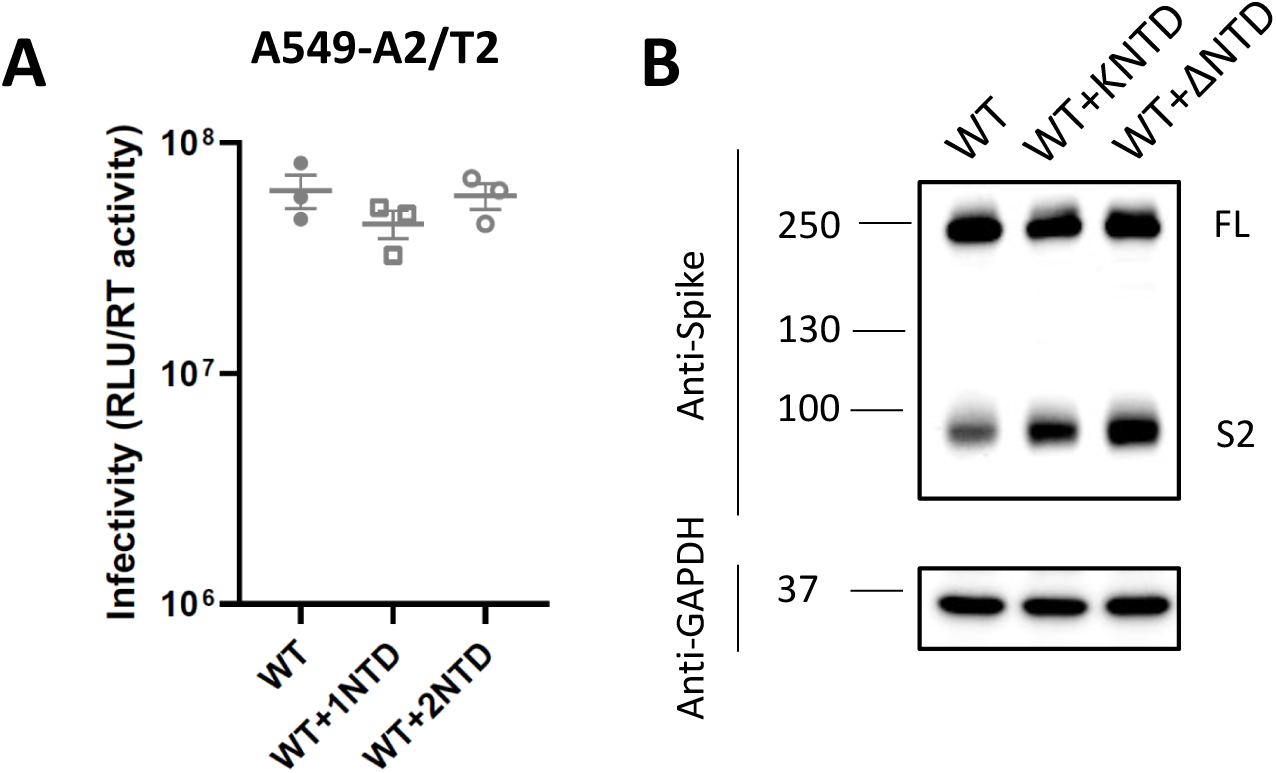
(A) The infectivity of WT, WT with Kappa NTD or Delta NTD chimeras in A549-A2/T2 cells (A) and western blot showing spike cleavage (B).

**Supplementary Figure 3:**
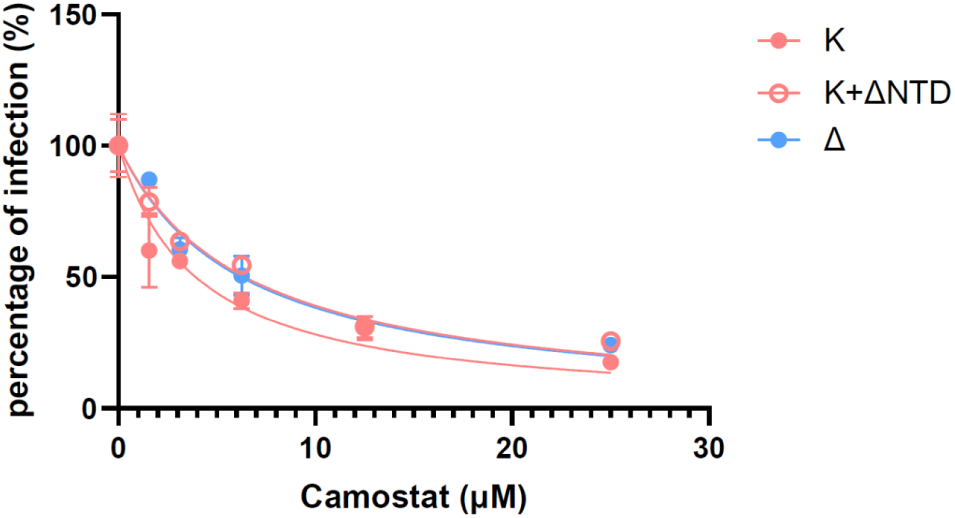
The Delta NTD fused with Kappa increases camostat resistance to a similar level as Delta.

